# The autophagic membrane tether ATG2A transfers lipids between membranes

**DOI:** 10.1101/555441

**Authors:** Shintaro Maeda, Chinatsu Otomo, Takanori Otomo

## Abstract

An enigmatic step in *de novo* formation of the autophagosome membrane compartment is the expansion of the precursor membrane phagophore, which requires the acquisition of lipids to serve as building blocks. Autophagy-related 2 (ATG2), the rod-shaped protein that tethers phosphatidylinositol 3-phosphate (PI3P)-enriched phagophores to the endoplasmic reticulum (ER), is suggested to be essential for phagophore expansion, but the underlying mechanism remains unclear. Here, we demonstrate that human ATG2A is a lipid-transferring protein. ATG2A can extract lipids from membrane vesicles and unload them to other vesicles. Lipid transfer by ATG2A is more efficient between its tethered vesicles than between untethered vesicles. The PI3P effectors WIPI4 and WIPI1 associate ATG2A stably to PI3P-containing vesicles, thereby facilitating ATG2A-mediated tethering and lipid transfer between PI3P-containing vesicles and PI3P-free vesicles. Based on these results, we propose that ATG2-mediated transfer of lipids from the ER to the phagophore enables phagophore expansion.

## Introduction

Autophagy is the bulk degradation-recycling process that plays crucial roles in the maintenance of cellular homeostasis in eukaryotes (Choi, Ryter, & Levine, 2013; Levine & Kroemer, 2019; Mizushima & Komatsu, 2011; Reggiori & Klionsky, 2013). Upon the induction of autophagy, a portion of the cytoplasm is sequestered within the double-membraned autophagosome compartment and transported to lysosomes (Lamb, Yoshimori, & Tooze, 2013; Mizushima, Yoshimori, & Ohsumi, 2011). Autophagosome biogenesis begins with the nucleation of the phagophore (also called the isolation membrane) adjacent to the endoplasmic reticulum (ER), followed by the expansion of the phagophore into a large cup-shaped double-membraned structure, resulting in the engulfment of bulk cytoplasmic constituents of various sizes ranging from protein molecules to organelles. Notably, the edge of the cup-shaped phagophore is associated with the ER (Hayashi-Nishino et al., 2009; Uemura et al., 2014; Yla-Anttila, Vihinen, Jokitalo, & Eskelinen, 2009), and this association is maintained during phagophore expansion (Graef, Friedman, Graham, Babu, & Nunnari, 2013; Suzuki, Akioka, Kondo-Kakuta, Yamamoto, & Ohsumi, 2013). This intimate spatial relationship has led to the hypothesis that the ER feeds the phagophore with lipids, enabling phagophore expansion (Tooze & Yoshimori, 2010). The phagophore is enriched with the lipid molecule phosphatidylinositol 3-phosphate (PI3P) (Cheng et al., 2014; Obara, Noda, Niimi, & Ohsumi, 2008), whose role is to recruit the PROPPIN family PI3P effectors Atg18/WD-repeat protein interacting with phosphoinositides (WIPIs) (Atg18 in yeast and WIPI1-4 in mammals) (Barth, Meiling-Wesse, Epple, & Thumm, 2001; Dove et al., 2004; Mercer, Gubas, & Tooze, 2018; Proikas-Cezanne, Takacs, Donnes, & Kohlbacher, 2015; Proikas-Cezanne et al., 2004) and the Atg18/WIPI-binding protein autophagy-related 2 (ATG2) (Atg2 in yeast and ATG2A/B in mammals) (Mizushima et al., 2011; Obara, Sekito, Niimi, & Ohsumi, 2008; Shintani, Suzuki, Kamada, Noda, & Ohsumi, 2001; Velikkakath, Nishimura, Oita, Ishihara, & Mizushima, 2012; Wang et al., 2001). Yeast studies have shown that both Atg18 and Atg2 localize exclusively to the phagophore edge and are required for phagophore expansion (Gomez-Sanchez et al., 2018; Graef et al., 2013; Suzuki et al., 2013), and mammalian studies have echoed the importance of ATG2A/B in phagophore expansion (Kishi-Itakura, Koyama-Honda, Itakura, & Mizushima, 2014; Tamura et al., 2017; Velikkakath et al., 2012). However, the mechanism through which these proteins enable membrane expansion remains unknown (Mizushima, 2018).

We recently characterized the structural and biochemical properties of the human ATG2A-WIPI4 complex (Chowdhury et al., 2018). The 1938 residue-long ATG2A is folded into a 20 nm-long, 30 Å-wide rod with both tips composed of Pfam database-registered conserved domains: a “Chorein_N” domain at the N-terminus of one tip (referred to as the N tip) and an “ATG2_CAD” domain in the middle of the sequence of the other tip (referred to as the CAD tip). Chorein is one of vacuolar protein sorting 13 (VPS13) family protein that function at contact sites between various organelles (Kumar et al., 2018; Lang, John Peter, Walter, & Kornmann, 2015; Murley & Nunnari, 2016; Park et al., 2016); as the name suggests, the Chorein_N domain is found at the N-terminus of VPS13 proteins (Pfisterer et al., 2014; Velayos-Baeza, Vettori, Copley, Dobson-Stone, & Monaco, 2004). The ATG2_CAD domain is unique to ATG2, and WIPI4 is tightly bound adjacent to the CAD tip (Fig. 1A). The yeast Atg2-Atg18 and the rat ATG2B-WIPI4 complexes exhibit similar overall shapes, suggesting that the structure is evolutionarily conserved and, therefore, likely important for the function of these complexes (Chowdhury et al., 2018; Zheng et al., 2017). Indeed, each tip of the ATG2A rod can bind to a membrane, and simultaneous membrane binding of both tips results in tethering the two membranes ~10-20 nm apart (Chowdhury et al., 2018), the typical distance observed at various organelle contact sites (Gatta & Levine, 2017). ATG2 proteins rely on packing defects in the membrane bilayer to effect membrane binding: Atg2 associates with giant unilamellar vesicles containing phosphatidylethanolamine (PE), a lipid molecule with a small head group that introduces packing defects (Gomez-Sanchez et al., 2018), as well as small unilamellar vesicles (SUVs) (Kotani, Kirisako, Koizumi, Ohsumi, & Nakatogawa, 2018), whose high curvatures create packing defects (Harayama & Riezman, 2018). ATG2A also associates tightly with SUVs but only weakly with large unilamellar vesicles (LUVs) (Chowdhury et al., 2018), whose lower curvatures create fewer packing defects. In correlation with the strength of the interactions, ATG2A/Atg2 tethers SUVs (Fig. 1B) (Chowdhury et al., 2018; Kotani et al., 2018), and ATG2A cannot tether LUVs (Fig. 1C) (Chowdhury et al., 2018). However, the ATG2A-WIPI4 complex can mediate homotypic tethering between two PI3P-containing LUVs as well as heterotypic tethering between a PI3P-containing LUV and a PI3P-free LUV (Fig. 1D) (Chowdhury et al., 2018), presumably because the N tip has affinity to LUVs and WIPI4 can direct the CAD tip to the PI3P-containing LUV. These data led us to propose that the ATG2A-WIPI4 complex tethers PI3P-positive phagophores to neighboring membranes, such as the ER and vesicles (Chowdhury et al., 2018). This model is supported by yeast studies concluding that the Atg2-Atg18 complex mediates the ER-phagophore association (Gomez-Sanchez et al., 2018; Kotani et al., 2018).

**Figure 1.**
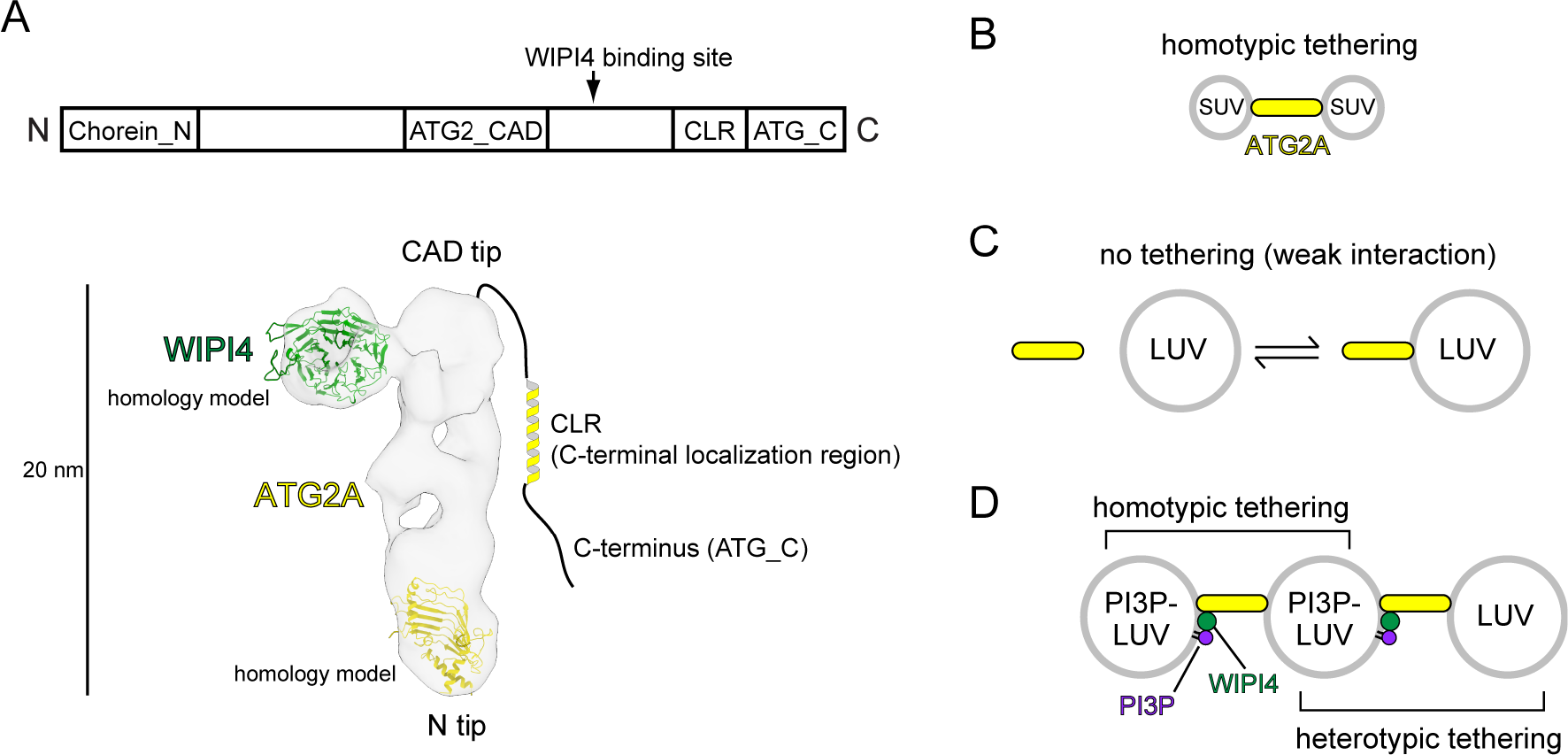
Structural and biochemical properties of the ATG2A-WIPI4 complex. (A) Summary of previous structural characterizations of the ATG2A-WIPI4 complex. The conserved and functional regions of ATG2 are indicated in the bar representation of the primary structure (top). The negative stain EM map (EMD-8899) of the ATG2A-WIPI4 complex is shown with docked homology models of WIPI4 and the ATG2A N-terminus as well as additional information regarding the C-terminal regions. The homology models were obtained from the I-TASSER server (Roy, Kucukural, & Zhang, 2010), and the molecular model was generated using ChimeraX (Goddard et al., 2018). (B-D) Summary of the membrane binding/tethering properties of ATG2A and the ATG2A-WIPI4 complex reported previously (Chowdhury et al., 2018). (B) ATG2A binds tightly to and tethers SUVs. (C) ATG2A associates weakly with and does not tether LUVs. (D) The ATG2A-WIPI4 complex can tether PI3P-containing LUVs and tether a PI3P-containing LUV to a PI3P-free LUV. Note that the tethering events shown in (B) and (C) lead to vesicle clustering in test tubes, but, for clarity, only one pair of tethered vesicles for each tethering pattern is shown.

ATG2 contains two additional conserved regions: an “ATG_C” domain, also registered in Pfam and located at the end of the C-terminus; and another short region preceding the ATG_C domain (Fig. 1A). Despite the name, ATG_C domains are also found in VPS13 proteins and are responsible for the localization of both ATG2A/B and VPS13 proteins to lipid droplets (Kumar et al., 2018; Tamura et al., 2017). Consistently, the ATG_C domain of ATG2A/B is dispensable for autophagy (Tamura et al., 2017). In contrast, the region preceding the ATG_C domain is required for the localization of ATG2A/B and Atg2 to phagophores (Kotani et al., 2018; Tamura et al., 2017; Velikkakath et al., 2012). This region, referred to as the C-terminal localization region (CLR), contains amphipathic α-helices that can associate with membranes (Chowdhury et al., 2018; Kotani et al., 2018; Tamura et al., 2017). The CLR of yeast Atg2 is required for the membrane tethering activity of this protein *in vitro* (Kotani et al., 2018) but is not required for that of ATG2A, consistent with the structural observation that the rod is the membrane tethering unit of ATG2A (Chowdhury et al., 2018). However, no rational explanations have been provided to clarify this discrepancy. Previous structural studies could not identify the location of the ATG2A/B C-terminus in the electron microscopy (EM) maps, suggesting that the C-terminus is flexible (Chowdhury et al., 2018; Zheng et al., 2017). Such flexibility may be advantageous for the C-terminal regions to function sufficiently as localization determinants (Kotani et al., 2018; Tamura et al., 2017; Velikkakath et al., 2012).

These recent advancements defined the role of ATG2-Atg18/WIPI4 complexes as mediators of the ER-phagophore edge association but did not explain the means by which such membrane associations lead to phagophore expansion. The size, overall shape, and membrane tethering activity of ATG2A called to mind tubular lipid-transferring proteins, such as the extended synaptotagmins and the ER-mitochondrial encounter structure (ERMES) complex (AhYoung et al., 2015; Jeong, Park, Jun, & Lee, 2017; Schauder et al., 2014; Wong, Gatta, & Levine, 2018), which further led us to hypothesize that ATG2A could be a lipid-transferring protein. This hypothesis is strongly supported by a recent study reporting that VPS13 family proteins are lipid-transferring proteins (Kumar et al., 2018). The crystal structure of the N-terminal fragment of yeast Vps13p revealed a unique structure with a large hydrophobic cavity that can apparently accommodate an unusually large number (~10) of glycerolipid molecules with a broad specificity and transfer those lipids between membranes. Homology modeling suggests that the structure of the ATG2 N-terminus would be very similar to that of Vps13p (Figs. 1A and S1). The VPS13 structure and the ATG2A structure proposed by the homology model fit into the N tip of our previous negative stain EM density map of ATG2A and occupy ~20-25% of the total volume (Fig. 1A) (Otomo, Chowdhury, & Lander, 2018). The central β-sheet, which serves as the base of this structure, is expected to extend in the C-terminal direction in the full-length protein (Fig. S1) (Kumar et al., 2018), which would create an elongated groove-shaped hydrophobic cavity in the ATG2A rod. Such a cavity could accommodate a large amount of lipids, as previously reported for VPS13 (Kumar et al., 2018).

Lipid transfer by ATG2 could be the fundamental molecular mechanism underlying phagophore expansion. Therefore, we investigated whether ATG2 is a lipid-transferring protein by performing a series of lipid transfer assays with ATG2A. Our strategy for this study was based on the membrane binding/tethering properties of ATG2A described above (Fig. 1B-1D). Our new results described below collectively show that ATG2A can indeed transfer lipids between membranes. WIPI4/1-enabled tethering facilitates ATG2A-mediated lipid transfer between a PI3P-containing membrane and a PI3P-free membrane. The demonstration of lipid transfer between membranes recapitulating a PI3P-positive phagophore and the ER allows us to propose that ATG2A-mediated lipid transfer drives phagophore expansion.

## Results

### ATG2A extracts lipids from membranes and unloads the lipids to membranes

Lipid transfer proteins can extract lipid molecules from existing membranes (Wong et al., 2018). To examine whether ATG2A has such activity, we adapted a lipid extraction assay in which liposomes containing fluorescent lipids (nitrobenzoxadiazole-conjugated PE: NBD-PE) are incubated with proteins, followed by liposome flotation to isolate the lipid-bound proteins (Kawano et al., 2018). To isolate ATG2A efficiently, we used LUVs prepared by extrusion through a filter with a pore size of 100 nm, because ATG2A would interact only weakly with those LUVs (Fig. 1C). After incubation of ATG2A with LUVs containing 20% NBD-PE and 80% dioleoylphosphatidylcholine (DOPC), the proteins and LUVs were separated by Nycodenz gradient-based liposome flotation (Fig. 2A). Upon centrifugation, the fluorescence signal of the protein-free control migrated from the bottom to the top of the tube, confirming that the LUVs floated (Fig. 2B and 2C). Similarly, the top fraction of the sample containing ATG2A became fluorescent, but the bottom remained so as well (Fig. 2B and 2C). SDS-PAGE analysis showed that most of the ATG2A proteins were in the bottom fraction (Fig. 2D), confirming that ATG2A did not bind tightly to these LUVs. Native PAGE analysis of the bottom fractions showed that the protein band was fluorescent (Fig. 2E), indicating that NBD-PE was bound to the protein. Quantification of the NBD fluorescence (Fig. 2C) and protein concentration (Fig. 2D) suggested that ATG2A was bound to ~6 molecules of NBD-PE. Thus, similar to VPS13 (Kumar et al., 2018), ATG2A can sequester a large amount of lipids. Given that NBD-PE constituted only 20% of the total lipids of the vesicles, it is likely that even more lipids are bound to the protein. Collectively, these data demonstrate that ATG2A can extract NBD-PE from membranes and dissociate from the membrane with those lipids attached.

**Figure 2.**
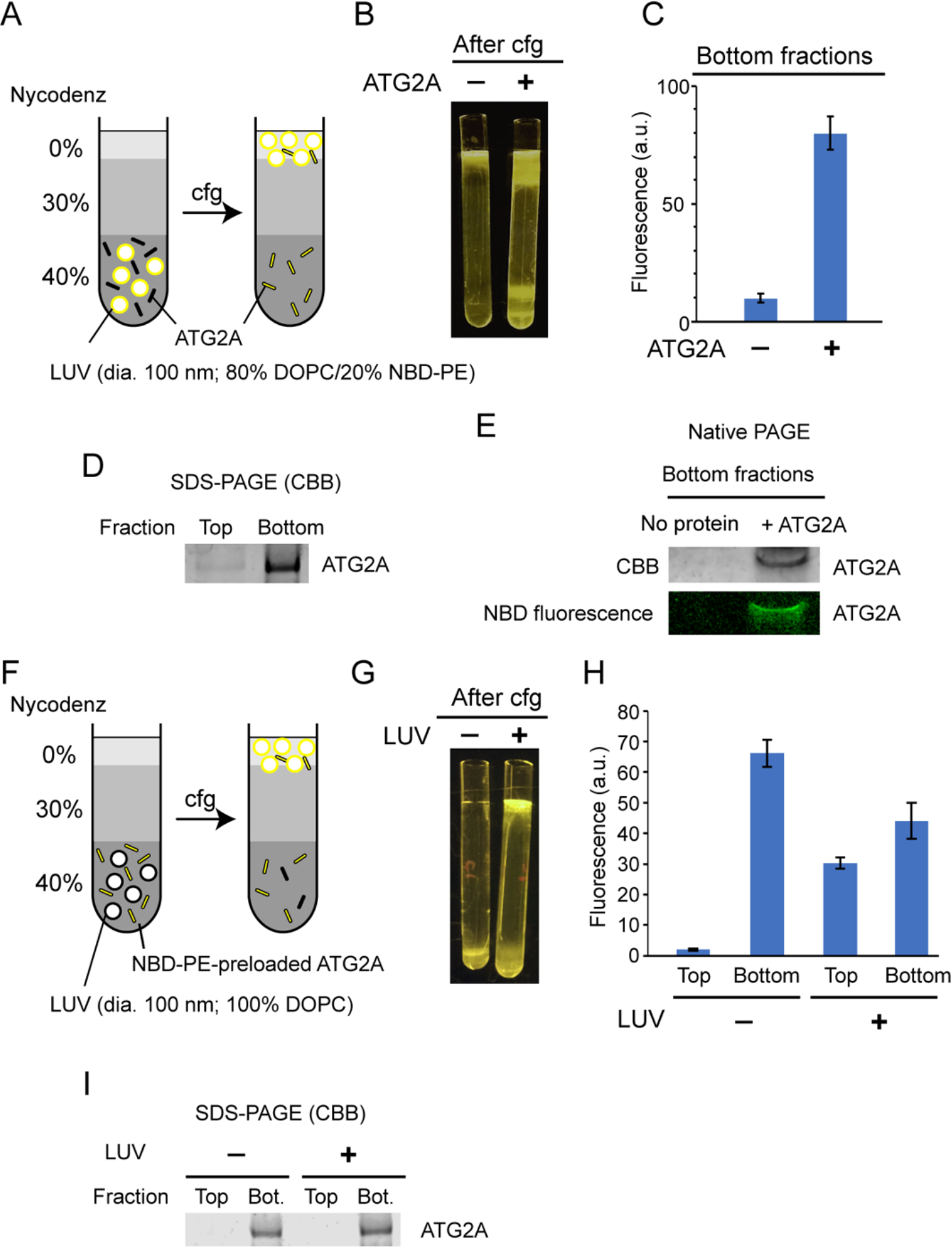
ATG2A extracts fluorescent lipids from membranes and unloads the lipids onto membranes. (A-E) NBD-PE extraction assay. (A) Diagram of the assay based on density (Nycodenz) gradient centrifugation. LUVs were prepared by extrusion using a 100 nm filter. (B) A postcentrifugation fluorescence image of the centrifuge tubes with and without ATG2A. (C) Quantification of the fluorescence signal in the bottom fractions of the tubes shown in (B). (D) SDS-PAGE with Coomassie blue (CBB) staining of the bottom fractions in (B). (E) Native PAGE of the bottom fractions in (B). The nonionic detergent decyl maltoside (0.2%) was added to the samples to prevent protein aggregation in the wells of the gel. CBB staining (top) and fluorescence (bottom) images are shown. (F-I) NBD-PE unloading assay. (F) Diagram of the assay. NBD-PE-preloaded ATG2A obtained from the bottom fraction in (B) was mixed with nonfluorescent LUVs (100 nm, 100% DOPC) and subjected to density gradient centrifugation. (G) A postcentrifugation fluorescence image of the centrifuge tubes with and without LUVs. (H) Fluorescence signal in the fractions in (G). (I) SDS-PAGE with CBB staining of the fractions in (G). Experiments were repeated three times. The fluorescence data in (C) and (H) and are shown as the average of the three repeats with the SD.

We next tested the unloading of fluorescent lipids from the protein. The bottom fractions from the extraction assay described above were collected and incubated with nonfluorescent LUVs (100% DOPC) or buffer as the control and were then resubjected to liposome flotation (Fig. 2F). After centrifugation, the fluorescence signal of the control remained in the bottom fraction, whereas that of the sample containing LUVs was detected not only in the bottom but also the top fraction (Fig. 2G and 2H). In both samples, the ATG2A proteins remained in the bottom fraction (Fig. 2I). These data indicate that ATG2A unloads NBD-PE onto LUVs during a transient interaction with the LUVs. The bottom fraction of the sample with LUVs yielded less (~33%) fluorescence signals than that of the LUV-free control. While this indicates that a substantial amount of lipids remained bound to the proteins, the signal reduction confirms the unloading of NBD-PE. In conclusion, the lipid extraction and unloading capabilities of ATG2A demonstrated here define ATG2A as a lipid-transferring protein.

### ATG2A transfers lipids between tethered membranes

The inefficient unloading observed above led us to hypothesize that tethering membranes by ATG2A could facilitate lipid transfer. We first examined whether lipids can be transferred between membranes tethered by ATG2A. To this end, we turned to our previous finding that ATG2A tethers SUVs efficiently (Fig. 1B) and performed a kinetic fluorescence lipid transfer assay with ATG2A and SUVs (Fig. 3A). In this accepted assay (Struck, Hoekstra, & Pagano, 1981), two kinds of vesicles are prepared and used as a mixture: donor vesicles that contain a pair of fluorescent lipids—NBD-PE and Rhodamine-PE (Rh-PE)—and acceptor vesicles that contain neither of the fluorescent lipids. Initially, NBD fluorescence arises solely from the donor vesicles and is suppressed due to quenching by Rhodamine on the same vesicle. However, owing to the dilution, NBD fluorescence increases upon the transfer of NBD-PE (or both NBD-PE and Rh-PE) to the acceptor vesicles. To stabilize the ATG2A-SUV association, we prepared donor and acceptor SUVs by sonication and included 25% dioleoylphosphatidylethanolamine (DOPE) and 25% dioleoylphosphatidylserine (DOPS) in both vesicles. The role of PE was described above, and DOPS improves the ATG2A-membrane association (Chowdhury et al., 2018). As shown in Fig. 3B, NBD fluorescence increased in the presence but not the absence of ATG2A proteins. A control experiment performed with no acceptor SUVs yielded no increase in NBD fluorescence. Thus, the signal increase observed in the presence of both SUVs resulted from the transfer of NBD-PE from the donor SUVs to the acceptor SUVs but not from the solubilization of NBD-PE by the proteins. Note that the increase in NBD fluorescence could also be explained by hemifusion or fusion between the donor and acceptor vesicles. However, these possibilities are ruled out by our previous data showing that ATG2A-clustered liposomes are disassembled upon protease treatment (Chowdhury et al., 2018). To further confirm this finding, we added sodium dithionite to the postreaction mixtures, which resulted in a decrease in NBD fluorescence in both samples—with and without protein—to the same level (~50% of the initial signal) (Fig. 3B). As explained in Fig. 3A, these results indicate that fusion did not occur.

**Figure 3.**
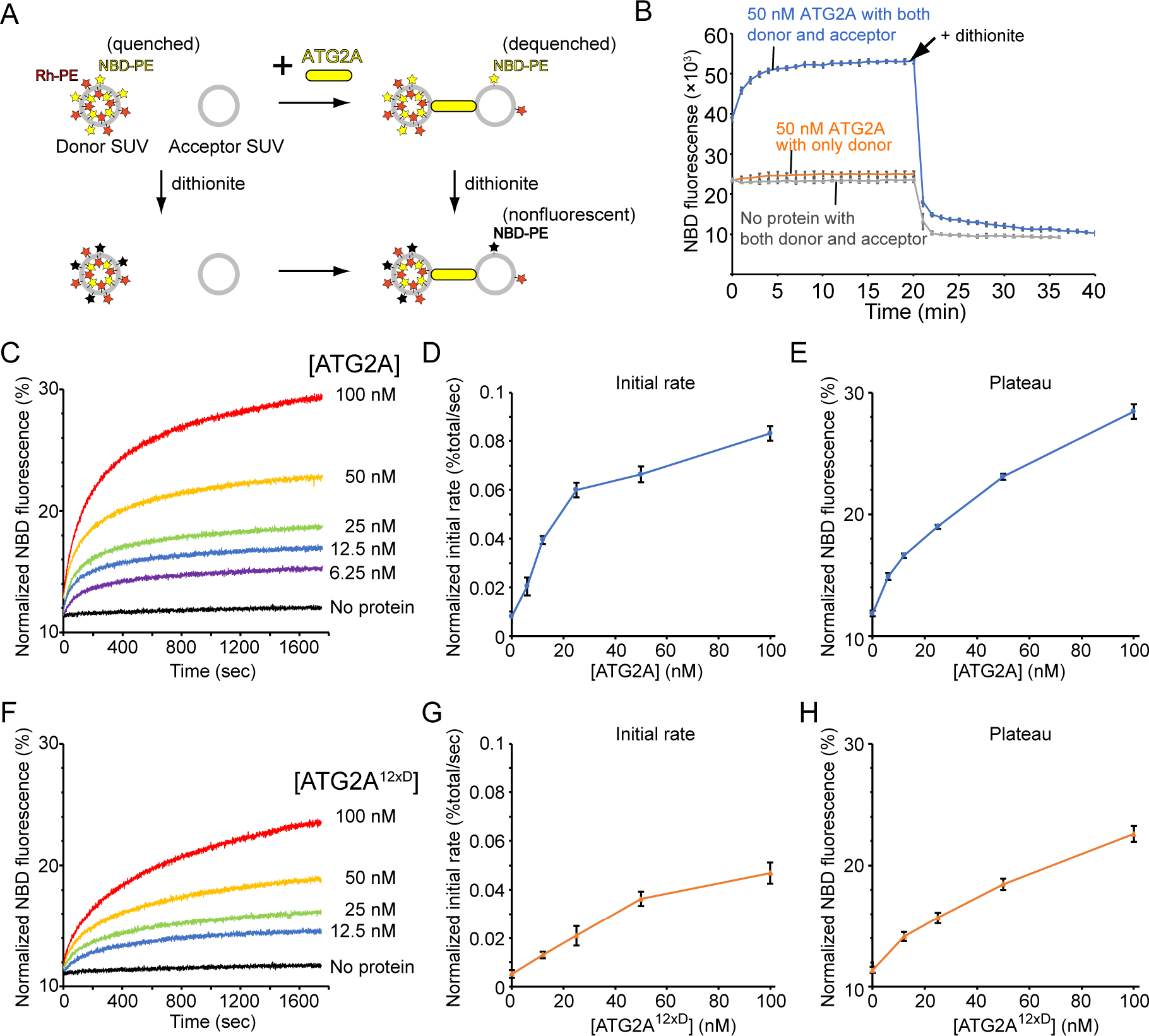
ATG2A transfers lipids between SUVs. (A) Diagram of the NBD fluorescence-based lipid transfer assay with ATG2A and SUVs. NBD fluorescence increases upon dilution of NBD, which is interpreted as the transfer of NBD-PE from the donor to the acceptor. Addition of dithionite would cause the loss of NBD fluorescence from the outer leaflets of liposomes (Meers, Ali, Erukulla, & Janoff, 2000). If the donor and the acceptor vesicles fuse, NBD-PE on the inner leaflet is diluted, thereby contributing to the signal increase. Hence, dithionite treatment increases the fluorescence of a membrane-fused sample to a higher level than that of a nonfused sample. (B) Lipid transfer assay with 50 nM ATG2A and 25 μM (lipid concentration) each of the donor SUVs (46% DOPC, 25% DOPE, 25% DOPS, 2% NBD-PE, and 2% Rh-PE) and the acceptor SUVs (50% DOPC, 25% DOPE, and 25% DOPS). SUVs were prepared by sonication, and the experiments were performed at 30°C. Each data point is presented as the average of the three independent experiments and shown with the SD. Sodium dithionite was added at the time indicated by the arrow. (C) Titration of the ATG2A protein in a lipid transfer assay with the same donor and acceptor vesicles as in (B). The experiments were performed at 25°C. The concentrations of ATG2A are indicated. NBD fluorescence was normalized to the maximum NBD fluorescence, which was measured upon the addition of 0.1% Triton X-100 at the end of each experiment. (D) Plot of the initial rate versus the concentration of ATG2A. (E) Plot of the plateau versus the concentration of ATG2A. (F-H) The same set of experiments (F) and analyses (G, H) as shown in (C-E) on ATG2A^12×D^.

The previous liposome flotation assays demonstrating the association of ATG2A with SUVs involved a 1.5-hr centrifugation step, allowing us to assume that most of the ATG2A proteins would remain associated with SUVs during the time course of the lipid transfer experiment. Under this assumption, the excess substrate SUVs in the reaction would be left unused, thus limiting the dynamic range of the signal (plateau) for a given concentration of ATG2A. We therefore performed a titration of ATG2A. Overall, the results showed that ATG2A facilitates lipid transfer in a concentration-dependent manner (Fig. 3C). For ATG2A concentrations of up to 20 nM, the rate increased linearly, confirming that the reaction was catalyzed by ATG2A (Fig. 3D). Beyond this concentration, the rate increase became nonlinear, indicating that the number of substrate SUVs became limiting. Moreover, the rates at these concentrations are underestimated due to overly fast reaction kinetics. For each time course, the signal first increased rapidly and then plateaus. Importantly, the plateau, defined as the signal at the final time point of each time course, increased as the protein concentration increased (Fig. 3E), indicating that lipids were transferred between tethered vesicles. Note that the plateau did not increase linearly with the protein concentration, further suggesting that the substrate SUVs were limiting. In addition, our initial assumption seems imperfect, because accurately, the signals continued to increase very gradually. This phenomenon suggests slow turnover of ATG2A, i.e., ATG2A occasionally dissociates from the donor or the acceptor, or even both, and acts on another vesicle, at least under these experimental conditions. These deviations from ideal kinetics, however, do not affect our conclusion that ATG2A can transfer NBD-PE between the tethered membranes.

We also tested a mutant protein, referred to as ATG2A^12×D^, in which twelve hydrophobic residues in the amphipathic helices in the CLR are mutated to aspartic acid (Chowdhury et al., 2018). Although CLR fragments containing these mutations cannot bind to membranes, and the mutations include a similar set of mutations that impair autophagy (Tamura et al., 2017), ATG2A^12×D^ is competent in membrane tethering *in vitro* (Chowdhury et al., 2018). As shown in Fig. 3F-3H, ATG2A^12×D^ exhibited lipid transfer activity similar to that of wild-type ATG2A, indicating that ATG2A^12×D^ transfers NBD-PE between the tethered membranes. Notably, the transfer rate and plateau are slower and lower, respectively, than those for the same concentration of wild-type ATG2A. The slower rate could indicate that the CLR contributes directly to lipid transfer. Given its affinity for membranes, the CLR may interact directly with the lipids being transferred, thereby affecting the rate of lipid transfer. However, such a role of the CLR would not explain the lower plateaus, which instead indicate that the membrane tethering activity of ATG2A^12×D^ is less potent than that of wild-type ATG2A. We previously reported that the 12×D mutation marginally altered the membrane tethering activity (Chowdhury et al., 2018). However, the dynamic light scattering and vesicle pull-down experiments utilized to monitor membrane tethering in the previous study are not very sensitive to a small decrease in the tethering activity of each molecule because tethering of freely diffusing liposomes leads to their clustering, which is facilitated by a multivalent effect involving multiple tethering molecules; therefore, the outputs of these experiments (the sizes of the clusters in the dynamic light scattering assays or the numbers of vesicles in the clusters in the vesicle pull-down assays) would not scale with the numbers of these molecules. In contrast, the kinetic lipid transfer assay should be more sensitive than these other assays, because lipid transfer is measured as the sum of all individual transfer events, each of which is catalyzed by one molecule; therefore, the output scales with the number of molecules. The contribution of the CLR to membrane tethering is consistent with the findings that the CLR fragment can bind to membrane vesicles through its amphipathic helical regions (Chowdhury et al., 2018) and that the corresponding region of yeast Atg2 is essential for the Atg2-membrane association (Kotani et al., 2018). Further work is required to clarify whether the CLR also plays a direct role in lipid transfer. Regardless of the precise role of the CLR, however, the presented results confirm that the rod region of the ATG2A protein is the core catalytic domain for both membrane tethering and lipid transfer.

### WIPI4/1 facilitates ATG2A-mediated lipid transfer

Next, we sought to directly test our hypothesis that membrane tethering facilitates lipid transfer. To compare lipid transfer between tethered membranes and nontethered membranes, we turned to our previous finding that the ATG2A-WIPI4 complex but not ATG2A alone can tether a PI3P-containing LUV to a PI3P-free LUV (Fig. 1D). The lipid transfer assay was adapted for this condition by replacing liposomes with LUVs, with the donors containing PI3P and the acceptors containing no PI3P (Fig. 4A). In the absence of WIPI4, ATG2A (100 nM) slightly increased NBD fluorescence (Fig. 4B), suggesting that ATG2A can transfer lipids between nontethered donors and acceptors but does so inefficiently. Strikingly, the addition of WIPI4 to ATG2A resulted in the acceleration of lipid transfer in a WIPI4 concentration-dependent manner, whereas the signal was not altered with WIPI4 alone (100 nM), demonstrating that WIPI4 facilitates ATG2A-mediated lipid transfer. Notably, the lipid transfer rate increased linearly with the concentration of WIPI4 and nearly saturated at the 1:1 stoichiometric concentration of WIPI4 to ATG2A (Fig. 4C), consistent with the strong interaction between WIPI4 and ATG2A. Based on these data, the above finding that ATG2A alone can transfer lipids between tethered SUVs, and the observation that the ATG2A-WIPI4 complex tethers PI3P-containing LUVs to PI3P-free LUVs (Chowdhury et al., 2018), we concluded that lipid transfer is facilitated by ATG2A-mediated membrane tethering.

**Figure 4.**
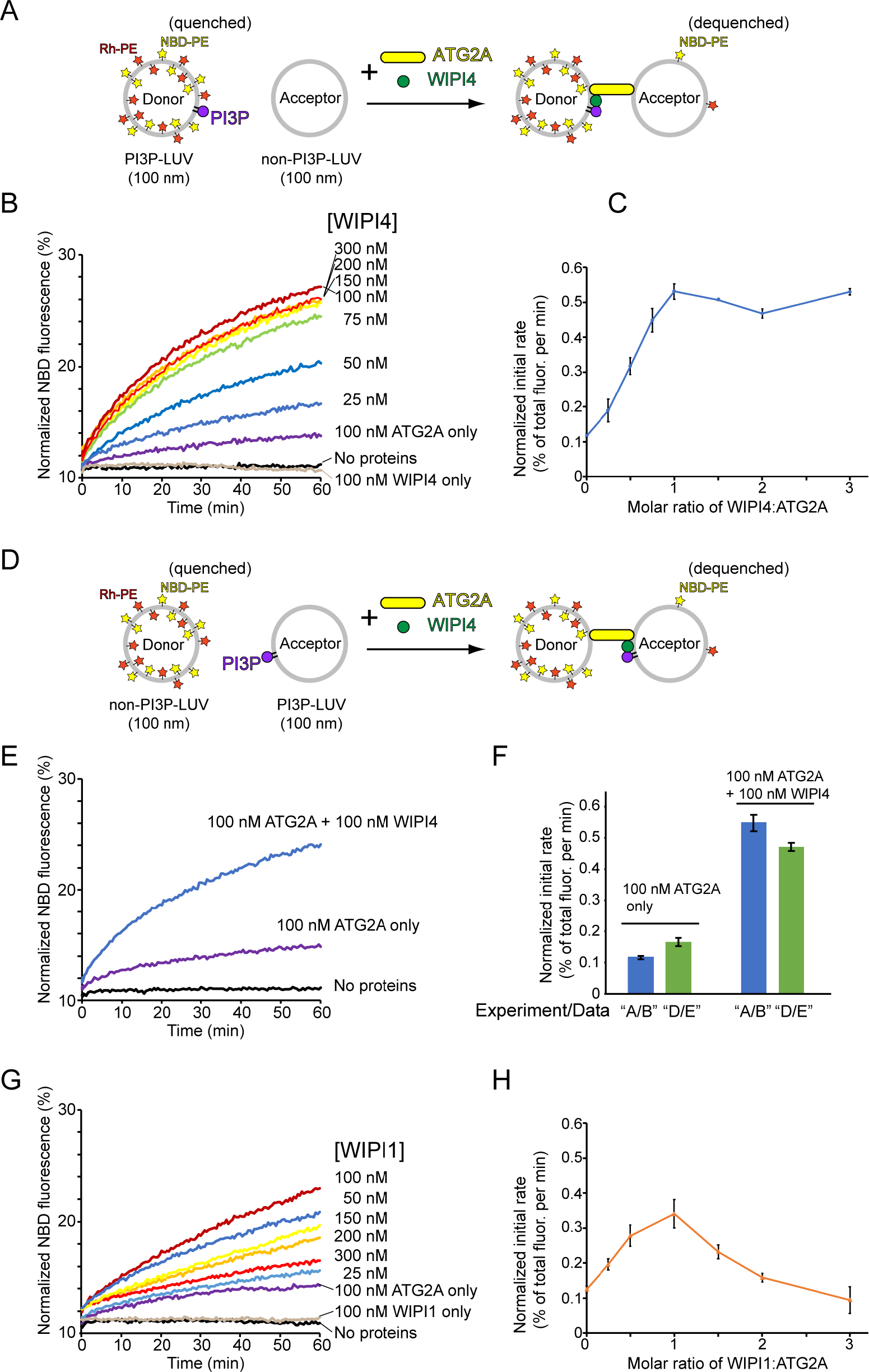
WIPI4 and WIPI1 facilitate ATG2A-mediated lipid transfer. (A) Diagram of the lipid transfer assay with PI3P-containing donor LUVs (5% PI3P, 46% DOPC, 25% DOPE, 20% DOPS, 2% NBD-PE, and 2% Rh-PE) and PI3P-free acceptor LUVs (50% DOPC, 25% DOPE, and 25% DOPS). (B) Titration of WIPI4 into 100 nM ATG2A. Protein concentrations are indicated. (C) Plot of the initial rates versus the molar ratios of WIPI4:ATG2A. (D, E) Diagram of (D) and data from (E) the lipid transfer assay with 25 μM PI3P-free donor LUVs (46% DOPC, 25% DOPE, 25% DOPS, 2% NBD-PE, and 2% Rh-PE) and PI3P-containing acceptor LUVs (5% PI3P, 50% DOPC, 25% DOPE, and 20% DOPS). Note that the position of PI3P is opposite from that in (A). (F) Comparisons of the initial rates of lipid transfer between the experiments performed in (A, B) and (D, E). Data obtained with 100 nM ATG2A in the presence or absence of 100 nM WIPI4 are compared. (G) Titration of WIPI1 into 100 nM ATG2A. The experiment was performed as in (A). (H) Initial rates in the experiment shown in (G). The experiments were performed at 25°C. For each experiment, the average of three repeats is shown with the SD.

In the experiment described above, PI3P was included in the donor vesicles, orienting the ATG2A-WIPI4 complex such that the WIPI4-CAD tip is bound to the donor and the N tip to the acceptor (Fig. 4A). NBD fluorescence increases mostly upon the transfer of NBD-PE from the donor to the acceptor vesicles, suggesting that ATG2A-mediated lipid transfer could be unidirectional. To examine this possibility, we reversed the orientation of the ATG2A-WIPI4 complex by including PI3P only in the acceptor vesicles (Figs. 4D). The lipid transfer experiments with this new pair of LUVs in the absence of WIPI or the presence of the stoichiometric concentration of WIPI4 showed lipid transfer rates similar to those found in the above experiment (Fig. 4E and 4F). Thus, ATG2A-mediated lipid transfer *in vitro* is an exchange reaction.

Four mammalian WIPI paralogs have been reported to function in different stages of autophagosome biogenesis (Proikas-Cezanne et al., 2015). The tight binding between WIPI4 and ATG2A has been advantageous for the structural and biochemical studies conducted thus far, but we wanted to determine whether other WIPIs can facilitate ATG2A-mediated lipid transfer. Among the other three WIPIs, we succeeded in obtaining only the WIPI1 protein with sufficient stability and quantity for these biochemical assays. As shown in Fig. 4G, ATG2A-mediated lipid transfer was accelerated in the presence of WIPI1. However, unlike with WIPI4, the increase in the rate occurred only up to the WIPI1:ATG2A stoichiometric concentration; further additions of WIPI1 beyond this concentration resulted in decreases in the rate (Fig. 4H). As described later, this suppression appears to be due to WIPI1-induced aggregation of PI3P-containing vesicles. Thus, we concluded that in principle, WIPI1 can also facilitate ATG2A-mediated lipid transfer.

### WIPI4/1 and ATG2A cooperatively associate with PI3P-containing membranes

The facilitation of ATG2A-mediated lipid transfer by the WIPI4 and WIPI1 proteins demonstrated above suggests that ATG2A is recruited to PI3P-containing membranes by these WIPI proteins. Although we previously did not examine the membrane binding of ATG2A in the presence of WIPI4, our observation that the ATG2A-WIPI4 complex mediated membrane tethering involving PI3P-containing LUVs implied that WIPI4 recruits ATG2A to the membrane (Chowdhury et al., 2018). Because the ability of WIPI1 to recruit ATG2A has not been reported, we investigated the membrane recruitment of ATG2A directly. To this end, we first examined the bimolecular interaction between WIPI1 and ATG2A using an affinity pull-down assay, which detected no binding of free WIPI1 to bead-immobilized ATG2A (Fig. 5A). To explain the ability of WIPI1 to facilitate ATG2A-mediated lipid transfer, we hypothesized that WIPI1 and ATG2A cooperatively associate with PI3P-containing membranes, as shown previously for yeast Atg18 and Atg2 (Kotani et al., 2018). To test this hypothesis, we performed liposome flotation assays with 5% PI3P-containing LUVs and the WIPI4/1 and ATG2A proteins. In the experiments with WIPI4 only, the top fractions contained WIPI4, but the amount of WIPI4 was much lower than that in the bottom fractions (Fig. 5B; lanes 4-9). In contrast, WIPI1 was mostly recovered from the top fractions (Fig. 5B; lanes 13-18). Thus, both WIPIs bind to these 5% PI3P-containing LUVs, but WIPI4 binds less strongly than WIPI1, consistent with the results of the previous study that demonstrated WIPI-membrane interactions by sedimentation assays (Baskaran, Ragusa, Boura, & Hurley, 2012). As expected, ATG2A alone did not bind stably to these LUVs (Fig. 5C; lanes 4-6). In stark contrast, in the presence of WIPI4, ATG2A was recovered almost completely from the top fractions (Fig. 5C; lanes 7-12), and WIPI4 recovery from the top fraction (~100% at 100 nM) was improved greatly compared to that in the WIPI4-only experiments (~20% at 100 nM) (Fig. 5B, lanes 4-9), indicating that WIPI4 and ATG2A cooperatively associate with PI3P-containing membranes.

**Figure 5.**
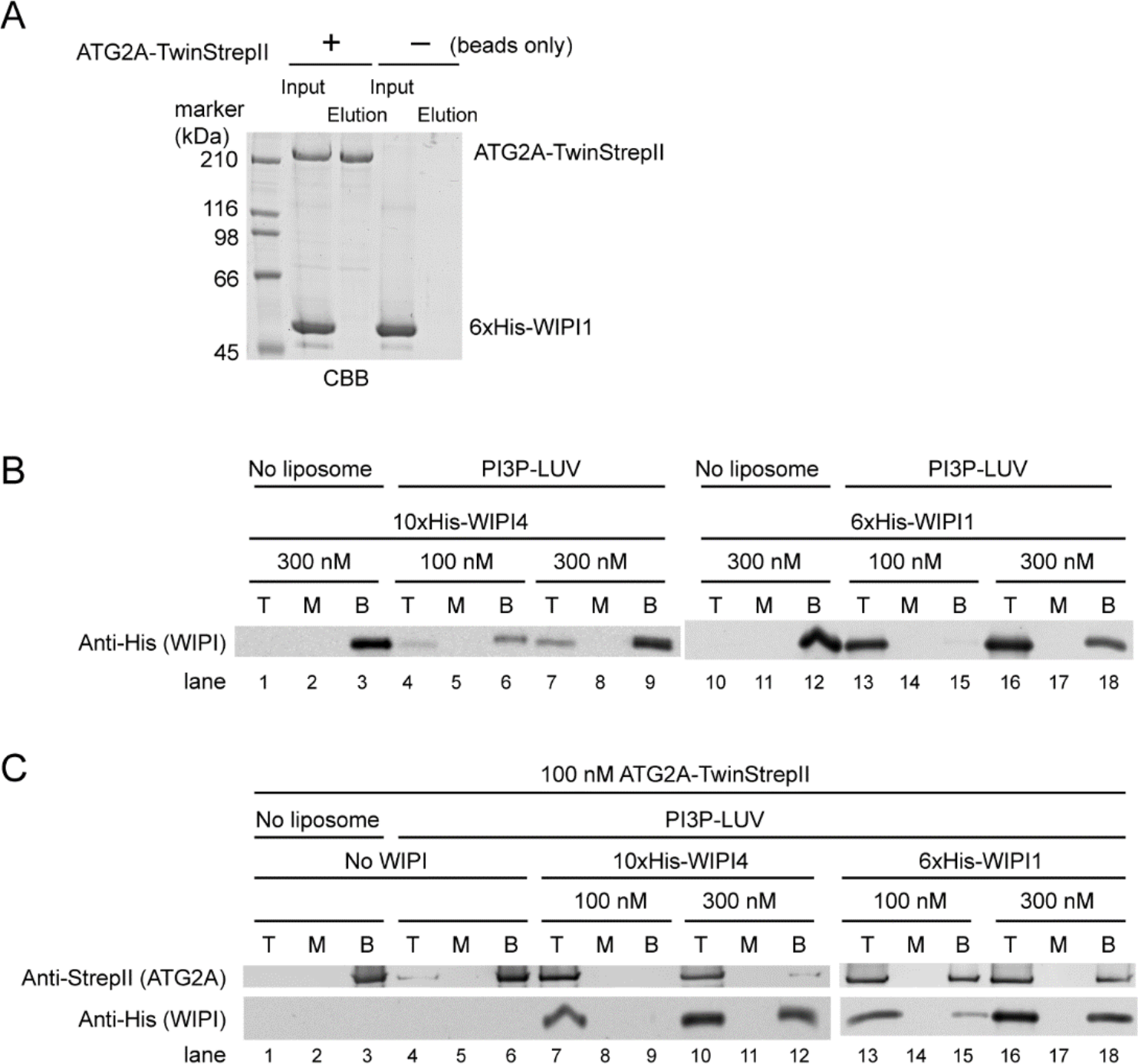
WIPI4 and WIPI1 recruit ATG2A to PI3P-containing LUVs. (A) Affinity pull-down experiments with ATG2A and WIPI1. Beads were loaded with ATG2A-TwinStrepII or no protein (beads only) and mixed with free WIPI1. Inputs and eluted fractions were analyzed by SDS-PAGE with CBB staining. (B) Liposome flotation assays to detect membrane binding of WIPI4 and WIPI1. LUVs (5% PI3P, 49% DOPC, 25% DOPE, 20% DOPS, and 1% DiD) were used at 25 μM. Protein concentrations are indicated. The top (T), middle (M), and bottom (B) fractions were collected after centrifugation and analyzed by western blotting. (C) Liposome flotation assays to examine ATG2A (100 nM) recruitment to PI3P-containing membranes by WIPI proteins. The experiments were performed as in (B).

The experiments with WIPI1 yielded similar results (Fig. 5C; lanes 13-18), demonstrating ATG2A recruitment to PI3P-containing membranes in the presence of WIPI1. Thus, WIPI1 appears to have an affinity for ATG2A that is too low to detect by the affinity pull-down assay but sufficient to synergize with the weak interaction between ATG2A and the PI3P-containing membranes for ATG2A recruitment. The percentages of ATG2A proteins recovered from the top fractions in the presence of WIPI1 (~60%) (Fig. 5C; lanes 13 and 16) were slightly lower than those in the presence of WIPI4 (~100%) (Fig. 5C; lanes 7 and 10), at least partially explaining the slower lipid transfer observed with WIPI1 than with WIPI4 at the stoichiometric concentration (100 nM) with ATG2A. In conclusion, in the presence of either WIPI protein, ATG2A binds stably to PI3P-containing membranes, providing the basis for WIPI-enabled membrane tethering and lipid transfer as demonstrated above.

### WIPI1 clusters PI3P-containing liposomes

In the experiments with WIPI1, we observed that the solutions containing 300 nM WIPI1 and PI3P-containing LUVs became slightly cloudy, which induced us to ask whether WIPI1 clusters those LUVs. To investigate this possibility, we performed turbidity assays with PI3P-containing LUVs and WIPI proteins and observed that the turbidity increased in a WIPI1 concentration-dependent manner (Fig. 6A), indicating vesicle clustering by WIPI1. The electron micrographs of the PI3P-containing LUVs with 300 nM WIPI1 show massive liposome aggregation, confirming the clustering (Fig. 6B). The liposome clustering was limited to WIPI1 because in the presence of WIPI4, the turbidity did not increase (Fig. 6A), and the liposomes were spread out in the EM images (Fig. 6B). The concentration dependence of WIPI1 clustering activity correlated with that of the suppression in lipid transfer shown above (Fig. 4G and 4H). Thus, WIPI1-induced aggregation of PI3P-containing donor LUVs would result in the sequestration of the proteins and donor LUVs from the PI3P-free acceptor LUVs, leading to the suppression of ATG2A-mediated lipid transfer at higher concentrations of WIPI1.

**Figure 6.**
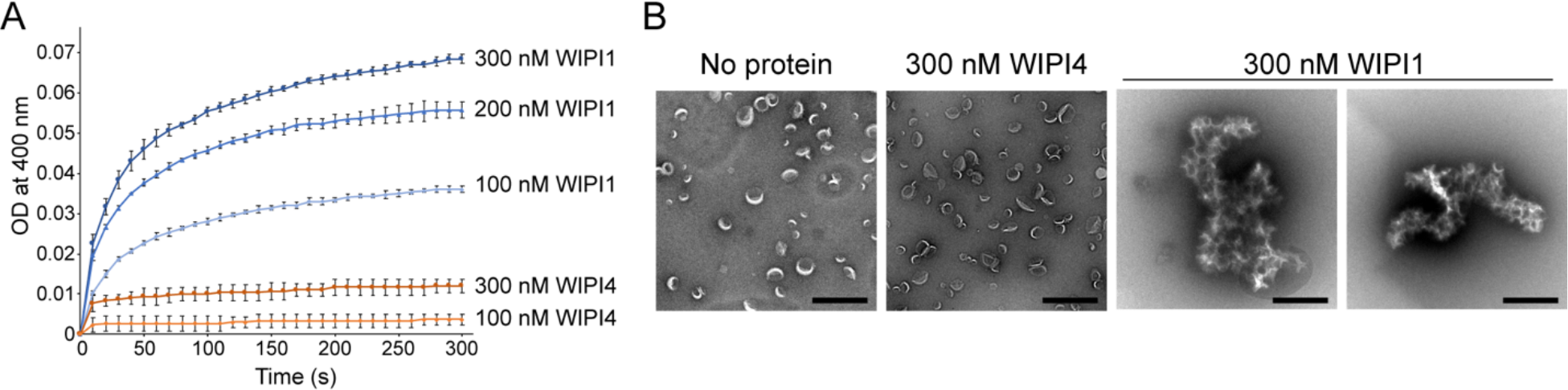
WIPI1 clusters PI3P-containing LUVs. (A) Turbidity assay. The OD at 400 nm of samples containing 25 μM LUVs (5% PI3P, 46% DOPC, 25% DOPE, and 20% DOPS) and WIPI4 or WIPI1 was measured. Protein concentrations are indicated. The SD were calculated from three independent experiments. (B) Micrographs of the negatively stained PI3P-LUVs in the presence and absence of 300 nM WIPI4 or WIPI1. Scale bar, 500 nm.

## Discussion

The results of yeast studies suggested that the Atg2-Atg18 complex directly tethers the phagophore to the ER (Gomez-Sanchez et al., 2018; Kotani et al., 2018). Atg18 is on the edge of PI3P-positive phagophores (Graef et al., 2013; Suzuki et al., 2013), directing the CAD tip of Atg2 to the phagophore and the N tip to the ER (Graef, 2018). ER anchoring by the N tip was demonstrated by Kotani *et al*., who created an autophagy-deficient Atg2 mutant by deleting the N-terminal 21 residues and restored autophagic activity by genetically fusing the transmembrane domain of the ER resident Sec71 protein to the N-terminus of the mutant (Kotani et al., 2018). In mammals, the precise locations of ATG2A/B on the expanding phagophore have not been established. However, WIPI4 has been shown to localize to the “omegasome” (Lu et al., 2011), the PI3P-positive transient region between the ER and the phagophore (Axe et al., 2008). Since WIPI4 and ATG2A bind cooperatively to PI3P-containing membranes (Fig. 5B), ATG2A is also likely recruited to the omegasome. According to previous electron tomography studies, omegasomes correspond to the thin tubular membranes that link the ER and the phagophore (Hayashi-Nishino et al., 2009; Uemura et al., 2014). Because this linker region and the phagophore are both PI3P-positive, it seems to be reasonable to consider that the omegasome represents the edge of the phagophore. The previous observations that the omegasome marker DFCP1 is localized adjacent to ATG proteins (Axe et al., 2008; Itakura & Mizushima, 2010) support this assumption. Furthermore, the representation of the phagophore edge by the PI3P-enriched omegasome is consistent with the PI3P-enrichment at the growing edge of the phagophore observed in yeast (Obara, Noda, et al., 2008). Thus, we now place the ATG2-WIPI complex between the ER and the phagophore edge, similar to the location suggested for the yeast Atg2-Atg18 complex (Kotani et al., 2018). Direct tethering of the ER and the phagophore edge by the ATG2-WIPI complex would allow the transfer of ER lipids to the phagophore, driving phagophore expansion (Fig. 7A).

**Figure 7.**
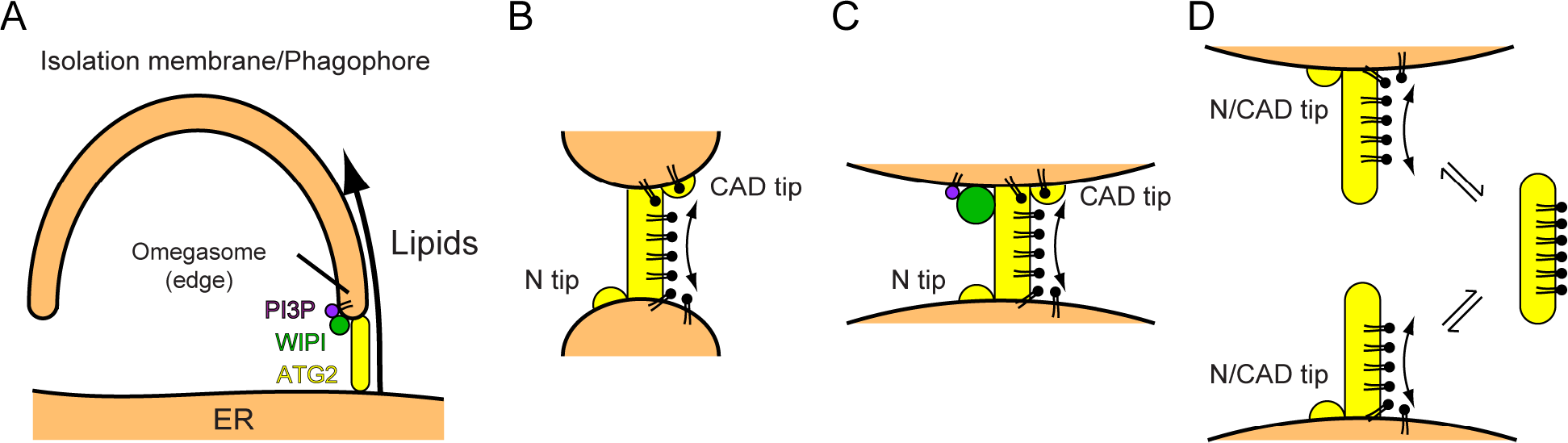
Models of phagophore expansion and ATG2A-mediated lipid transfer. (A) Illustration of ATG2-mediated transfer of lipids from the ER to the phagophore. (B, C) The bridge model for ATG2-mediated lipid transfer between highly-curved membranes (SUVs) in the absence of WIPI (B) and between less-curved membranes (LUVs) in the presence of WIPI (C). (D) A model of lipid loading/unloading in the absence of membrane tethering.

This model of phagophore expansion explains why and how the edge of the phagophore remains associated with the ER during its expansion (Graef et al., 2013; Suzuki et al., 2013) and why autophagosome membranes are of thinner types like the ER membrane (Arstila & Trump, 1968) but are devoid of ER proteins/markers such as cytochrome P-450, Dpm1, and GFP-HDEL (Suzuki et al., 2013; Yamamoto, Masaki, Fukui, & Tashiro, 1990). The recent observation that the ER-staining lipophilic dye octadecyl rhodamine B also stains the phagophore in yeast (Hirata, Ohya, & Suzuki, 2017) can be explained by the ATG2-mediated transfer of this lipophilic dye from the ER to the phagophore. Thus, phagophore expansion based on ATG2-mediated lipid transfer agrees with many previous lines of evidence. The high lipid binding capacity and membrane tethering-facilitated lipid transfer activity of ATG2A would enable this protein to transfer a large amount of lipids rapidly, an advantage for *de novo* membrane construction. Indeed, autophagosome formation is completed in ~5-10 min (Fujita et al., 2008; Geng, Baba, Nair, & Klionsky, 2008; Mizushima et al., 2001), and ATG2A-mediated lipid transfer between SUVs *in vitro* is mostly completed in a similar time frame (Fig. 3B). Notably, at least the edge of the phagophore is highly curved with a diameter of ~30 nm (Nguyen, Shteyn, & Melia, 2017), similar to the characteristics of SUVs. Thus, in principle, lipid transfer by ATG2A can account for the kinetics of phagophore expansion. An important implication that arises from this model is that to drive phagophore expansion, the equilibrium of ATG2A-mediated lipid transfer must be shifted toward the phagophore. Currently, no mechanism that allows such directionality for phagophore expansion is known. In addition, the type of energy that would be consumed in this process *in vivo* remains an important unknown.

The finding that WIPI proteins facilitate ATG2A-mediated lipid transfer suggests that PI3P dynamics control phagophore expansion: the number of PI3P molecules on the phagophore would determine the number of ATG2 proteins, thereby regulating the overall rate of phagophore growth. A reduction in the PI3P level, which occurs toward the end of autophagosome formation (Axe et al., 2008), would release ATG2 from the point of ER-phagophore contact, leading to the disruption of the contact and the cessation of phagophore expansion. This model of the regulation of phagophore expansion is supported by the previous observation that the overexpression of the PI3P-phosphatase myotubularin-related phosphatase 3 generates significantly smaller autophagosomes (Taguchi-Atarashi et al., 2010). According to this model, the PI3P effectors Atg18/WIPIs play a pivotal role in correlating PI3P dynamics to phagophore expansion. Atg18 partners primarily with Atg2, suggesting its linear relationship with phagophore expansion. In contrast, WIPI proteins, as a whole, play a broader role. For example, WIPI1/2 regulate the conjugation of LC3 family ubiquitin-like proteins to autophagosomal membranes (Bakula et al., 2017; H. E. Polson et al., 2010; H. E. J. Polson, Girardin, Wilson, & Tooze, 2014), and WIPI3 recruits the autophagosome initiator FIP200 (Bakula et al., 2017). Consistent with these ATG2A/B-independent functions, WIPI1/2 localize to the phagophore independently of ATG2A/B (Tamura et al., 2017; Velikkakath et al., 2012), a behavior also consistent with the strong associations of these proteins with PI3P-containing membranes (Baskaran et al., 2012). Furthermore, WIPI1 localizes not only to the edge of but also throughout the phagophore surface (Itakura & Mizushima, 2010). This distribution of WIPI1 across the phagophore is similar to that of LC3, supporting a role of WIPI1 in LC3 lipidation. The direct link between WIPIs and ATG2A/B has been limited to WIPI4, but our work adds WIPI1 to this link. Despite the failure to detect its interaction with ATG2A (Fig. 5A), WIPI1 can recruit ATG2A to PI3P-containing membranes and facilitate ATG2A-mediated lipid transfer (Fig. 5B). These findings raise the important possibility that the yet-untested WIPI2 and WIPI3, which, like WIPI1, do not interact strongly with ATG2A/B (Bakula et al., 2017), might also cooperate with ATG2A. The high sequence identity (~80%) between WIPI1 and WIPI2 (H. E. Polson et al., 2010) supports this potential role of WIPI2.

A notable difference between WIPI4 and WIPI1 in our study is the suppression of ATG2A-mediated lipid transfer that occurs at high concentrations of WIPI1 (Fig. 4G and 4H). While we suggested above that WIPI1-induced aggregation of PI3P-containing membranes would cause this suppression, we also note here that freely diffusible LUVs are not perfect models of the phagophore, which is a large double-membraned sac scaffolded by the ER that does not cluster unassisted. Therefore, we consider clustering-induced suppression to be an inevitable consequence of the LUV-utilizing *in vitro* experiment. Liposome clustering was likely induced by self-oligomerization of WIPI1, as has been reported for Atg18/WIPIs on membranes and in solution (Baskaran et al., 2012; Gopaldass, Fauvet, Lashuel, Roux, & Mayer, 2017; Scacioc et al., 2017). Interestingly, the self-oligomerization of Atg18 on giant unilamellar vesicles tubulates the membrane (Scacioc et al., 2017), and PI3P-positive thin tubular membranes at the phagophore edge (omegasomes) form clusters (Uemura et al., 2014). Thus, WIPI1 may mediate the clustering of these membranes.

The coupling between the membrane tethering and lipid transferring activities of ATG2A suggests a “bridge” model of lipid transfer (Fig. 7B and 7C). In this model, two membranes are stably tethered by ATG2, and lipids are transferred between the apposed membranes via ATG2. For ATG2 to serve as the path for lipid transfer, its hydrophobic cavity must be assumed to be extended almost completely from tip to tip so that lipids can slide through. Structural predictions propose such an elongated cavity for VPS13 (Kumar et al., 2018), consistent with the finding that VPS13 and ATG2A can bind a large amount of lipids. A probable mechanism for lipid loading/unloading emerges from the attempt to provide a structural explanation for Kotani *et al.*’s data showing that the N-terminal 46 residues of Atg2 can localize to the ER in isolation and that the deletion of the N-terminal 21 residues impairs the membrane tethering activity of Atg2 *in vitro* (Kotani et al., 2018). Those highly conserved residues at the N-terminus form an amphipathic helix with the hydrophobic side facing the cavity (Fig. S2). A hinge-like conformational change in the loop succeeding the N-terminal helix could flip out the helix and allow it to dock onto a membrane surface. Concomitantly, the cavity would be opened and docked to the membrane, allowing lipids to flow in. Such role of the N-terminal helix as the membrane anchor is also supported by the aforementioned data that the N-terminal 21 residues can be replaced by a transmembrane domain (Kotani et al., 2018). No structural information regarding the CAD tip is available, but we speculate that a similar mechanism might apply to this opposing tip because lipid transfer is facilitated by membrane tethering involving the CAD tip. Moreover, the bridge mechanism holds in the presence of WIPI proteins (Fig. 7C). Because they are adjacent to the CAD tip, WIPI4/1 would stabilize the CAD tip-membrane association, thereby enabling the CAD tip to participate in lipid transfer. While the role of WIPI4/1 in recruiting ATG2A to the PI3P-containing membrane sufficiently explains the data presented above, WIPI4/1 may also activate the CAD tip, which may be essential when the surface of the substrate membrane has few defects. The bridge model can explain why ATG2A can extract lipids from membranes with few surface defects but inefficiently unloads lipids onto those membranes: in a transient interaction with those membranes, ATG2A extracts lipids via only one of its two tips. The loading of lipids into an empty cavity from a membrane-bound tip would be energetically favored, but unloading from the same tip might not be, because lipids in the cavity are bound to the protein and would diffuse only slowly out of a cavity in which the opposite end was closed (Fig. 7D). Given that the N tip would have a higher affinity to LUVs than the CAD tip, loading from the N tip could be more probable than loading from the CAD tip in the nontethered situation. However, the CAD tip does have an affinity for LUVs, as evidenced by the cooperative binding of ATG2A and WIPI4/1 to PI3P-containing LUVs, which must be contributed by the CAD tip. When tethered, both tips are open to membranes, allowing lipids to diffuse more rapidly onto the ATG2 bridge (Fig. 7B and 7C). Future studies toward the elucidation of the mechanism of ATG2-mediated lipid transfer will build upon the bridge model described here.

## Acknowledgments

We thank Kazuto Ohashi for his contribution to preliminary studies for this project. We are grateful to Ashok Deniz for the generous advice and critical reading of the manuscript. This research was supported by a grant from the National Institute of General Medical Sciences (R01-GM092740).

## Materials and Methods

### Protein expression and preparation

Human ATG2A and WIPI4 were expressed and purified as described previously (Chowdhury et al., 2018). Human WIPI1 was expressed and purified using the same procedure used for WIPI4. ATG2A was genetically fused to an N-terminal TEV protease-cleavable glutathione S-transferase (GST) and to a C-terminal PreScission protease-cleavable TwinStrepII (TS) tag. The GST tag but not the TwinStrepII tag was removed during the purification. Purified ATG2A-TS was dialyzed against 20 mM HEPES (pH 7.5), 150 mM NaCl, and 1 mM Tris(2-carboxyethyl)phosphine (TCEP) and used for assays. WIPI4 and WIPI1 were expressed with N-terminal TEV protease-cleavable ten-histidine (10×His) and six-histidine (6×His) tags, respectively. WIPI4 and WIPI1 proteins from which the His tags were removed were used in the lipid transfer assays, and those with uncleaved His tags (10×His-WIPI4 and 6×His-WIPI1) were used in the liposome flotation assays.

### NBD-PE lipid extraction assay

Lipids (Avanti Polar Lipids) were mixed in a glass tube at a molar ratio of 80% 1,2-dioleoyl-*sn*-glycero-3-phosphocholine (DOPC) and 20% 1,2-dioleoyl-sn-glycero-3-phosphoethanolamine-N-(7-nitro-2-1,3-benzoxadiazol-4-yl) (NBD-PE) dissolved in chloroform and dried under a stream of nitrogen gas. The obtained lipid film was further dried under a vacuum for 30 min and then hydrated in 20 mM HEPES (pH 7.5) and 150 mM NaCl. The hydrated lipids were subjected to seven freeze-thaw cycles using liquid nitrogen and a water bath set to 42°C. The resulting LUV solution was passed more than 20 times through a polycarbonate filter membrane with a pore size of 100 nm using the extruder from Avanti Polar Lipids, Inc. The prepared LUVs were stored at 4°C in the dark. A 200 nM concentration of ATG2A protein was mixed with 100 nM LUVs in 20 mM HEPES (pH 7.5), 150 mM NaCl, and 1 mM TCEP (total volume, 37.5 μL). After incubation for 1 hr at ~22°C, an equal volume of 80% Nycodenz (Accurate Chemical) solution was added to the protein-LUV solution and mixed thoroughly. Then, the mixture was placed at the bottom of a centrifuge tube. A total of 475 μL of 30% Nycodenz solution was placed above the bottom layer of solution, and 50 μL of buffer with no Nycodenz (20 mM Hepes (pH 7.5), 150 mM NaCl, and 1 mM TCEP) was then placed on the top. The tubes were centrifuged in an SW55Ti rotor (Beckman Coulter) at 279,982 × *g* (max) for 1.5 hr at 18°C. After centrifugation, the top 520 μL was removed, and the bottom fraction (80 μL) was then collected and subjected to SDS-PAGE. The gels were scanned on a Typhoon 9410 imager (GE Healthcare) to quantify NBD fluorescence, followed by CBB staining and scanning on an Odyssey imager (LI-COR Biosciences) for protein visualization and quantification of the protein concentration. The bottom fractions were also subjected to native PAGE. Decyl maltoside was added to the sample fractions at a final concentration of 0.2% to prevent aggregation of the ATG2A protein in the wells of the gel. The gel was scanned on the Typhoon 9410 imager to detect NBD fluorescence, followed by CBB staining to detect proteins.

### NBD-PE lipid unloading assay

LUVs containing 100% DOPC were prepared exactly as described above. ATG2A loaded with NBD-PE was collected from the bottom fraction of the extraction assay described above, and DOPC-containing LUVs were added to this fraction at a final concentration of 100 nM. After a 1-hr incubation, 80% Nycodenz solution was added to the mixture to adjust the Nycodenz concentration to 40%. This solution (75 μL) was placed at the bottom of a centrifuge tube, and 30% Nycodenz (475 μL) and 0% Nycodenz (50 μL) were sequentially placed above the bottom solution. The tube was centrifuged as described above. The top and bottom fractions (80 μL each) were collected and subjected to SDS-PAGE analysis. The gels were scanned on Typhoon 9410 imager for quantification of fluorescence, followed by CBB staining to visualize proteins.

### Lipid transfer assay

The following liposomes were prepared as described above: donor SUVs—46% DOPC, 25% 1,2-dioleoyl-*sn*-glycero-3-phosphoethanolamine (DOPE), 25% 1,2-dioleoyl-*sn-*glycero-3-phospho-L-serine (DOPS), 2% NBD-PE, and 2% 1,2-dioleoyl-*sn*-glycero-3-phosphoethanolamine-N-(lissamine rhodamine B sulfonyl) (Rh-PE); acceptor SUVs—50% DOPC, 25% DOPE, and 25% DOPS; PI3P-containing donor LUVs—5% PI3P, 46% DOPC, 25% DOPE, 20% DOPS, 2% NBD-PE, and 2% Rh-PE; PI3P-free acceptor LUVs—50% DOPC, 25% DOPE, and 25% DOPS; PI3P-free donor LUVs—46% DOPC, 25% DOPE, 25% DOPS, 2% NBD-PE, and 2% Rh-PE; and PI3P-containing acceptor LUVs—5% PI3P, 50% DOPC, 25% DOPE, and 20% DOPS. Proteins were mixed with 25 μM (concentration of lipids) donor and acceptor vesicles in a cuvette and placed in a fluorimeter (Photon Technology International) at 30°C (only for the experiment shown in Fig. 3B) or 25°C. NBD fluorescence was recorded at an excitation wavelength of 460 nm and an emission wavelength of 535 nm every 1 second (Fig. 3) or 30 seconds (Fig. 4). At the end of each reaction, Triton X-100 was added to the reaction mixture to a final concentration of 0.1% (v/v) to solubilize all lipids and therefore maximize NBD fluorescence. All data points in each time course were normalized to the maximum fluorescence value. For the dithionite experiments, sodium dithionite dissolved in 50 mM Tris (pH 10) was added to the postreaction mixture at a final concentration of 5 mM. Rates were obtained by linear regression of the initial points in the time course data (5-25 seconds for Fig. 3 and 30-300 seconds for Fig. 4). Plateaus were defined as the normalized NBD fluorescence signals at the final time point of each time course.

### Affinity pull-down assay

ATG2A-TS was loaded onto Strep-Tactin Superflow beads in buffer containing 20 mM HEPES (pH 7.5), 150 mM NaCl, 0.5 mM TCEP, 20% glycerol, and 0.1% Triton X-100. After loading, the beads were rigorously washed with buffer and were then mixed with His-WIPI1. After ~5 min of incubation, the tube was centrifuged to collect the beads. The beads were washed three times, after which proteins were eluted in the same buffer supplemented with 5 mM desthiobiotin. The input and eluate solutions were subjected to SDS-PAGE.

### Liposome flotation-based binding assay

Liposome flotation assays were performed as described previously (Chowdhury et al., 2018). Briefly, ATG2A, WIPI4, and WIPI1 proteins were mixed with 25 μM LUVs (5% PI3P, 49% DOPC, 25% DOPE, 20% DOPS, and 1% 1,1’-Dioctadecyl-3,3,3’,3’-tetramethylindodicarbocyanine perchlorate (DiD) (Marker Gene Technologies)) as indicated in Fig. 5B and 5C in 150 μL of buffer containing 20 mM HEPES (pH 7.5), 150 mM NaCl, 1 mM TCEP, and 40% Nycodenz at the bottom of centrifugation tubes. A layer of 400 μL of 30% Nycodenz buffer was placed on the top of each sample, and a layer of 50 μL of 0% Nycodenz buffer was then placed on top. Tubes were centrifuged as described above. The top (150 μL), middle (300 μL), and bottom (150 μL) fractions were collected from top and analyzed by western blotting using anti-StrepII and anti-His tag antibodies.

### Turbidity assay

A 25 μM concentration of LUVs (5% PI3P, 50% DOPC, 25% DOPE, 20% DOPS) was mixed with 100, 200, or 300 nM WIPI1 or WIPI4 in 20 mM HEPES (pH 7.5), 150 mM NaCl, and 1 mM TCEP and incubated at ~22°C in a 50 μL cuvette placed in an Ultraspec 2000 UV spectrophotometer (Pharmacia Biotech). The optical density (OD) at 400 nm was recorded every 10 seconds.

### Negative stain electron microscopy

LUVs (100 μM) were mixed with 300 nM WIPI4 or WIPI1. After incubation for 5 min at ~22°C, a 3 μL drop of the mixture was placed on a glow discharge continuous carbon grid (Electron Microscopy Science) and stained with 2% uranyl acetate. Grids were imaged at a magnification of 11,500 × on an FEI Tecnai F20 transmission electron microscope operating at 200 keV.

---

**Figure S1.**
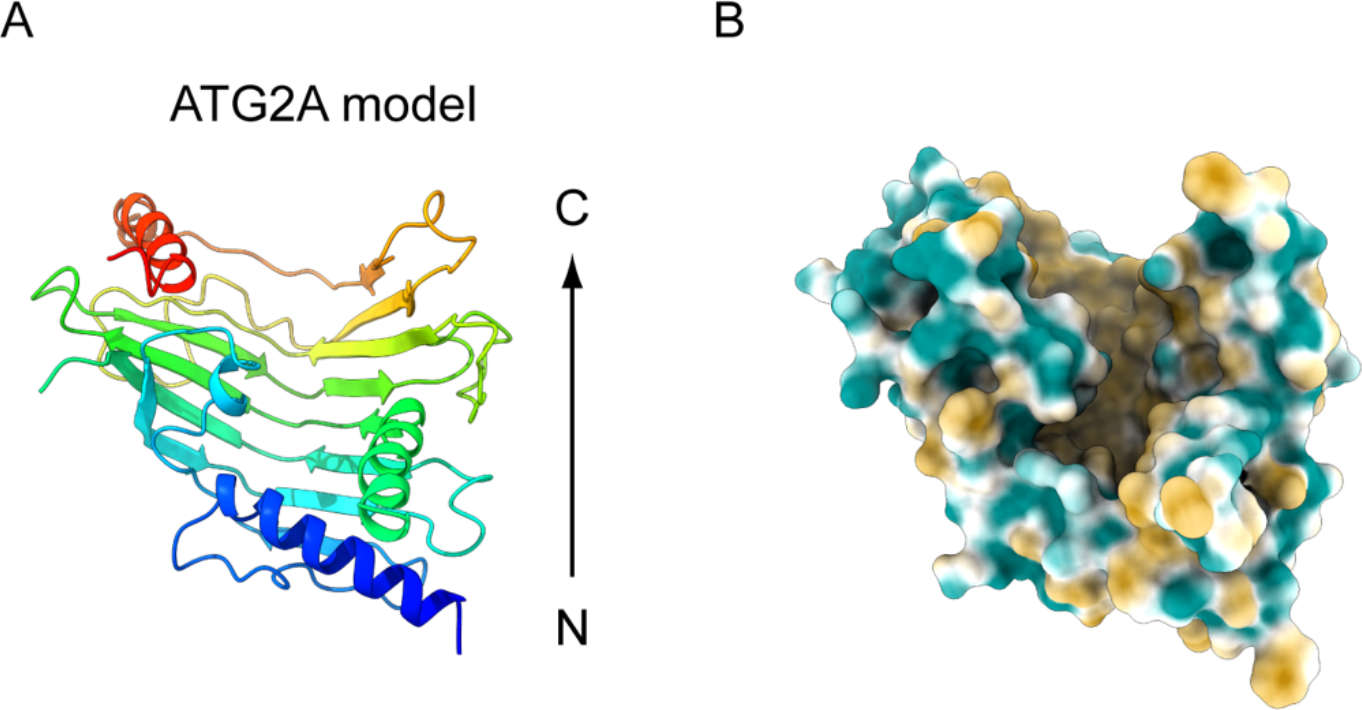
A homology model of the ATG2A N-terminus. (A) A cartoon representation of the ATG2A homology model built by the I-TASSER server based on the crystal structure of *Chaetomium thermophilum* Vps13p (PDB ID: 6cbc). The model is rainbow-colored from the N-terminus (blue) to the C-terminus (red). (B) A surface representation colored according to the hydrophobicity potential. Hydrophobic and hydrophilic residues are shown in khaki and turquoise, respectively.

**Figure S2.**
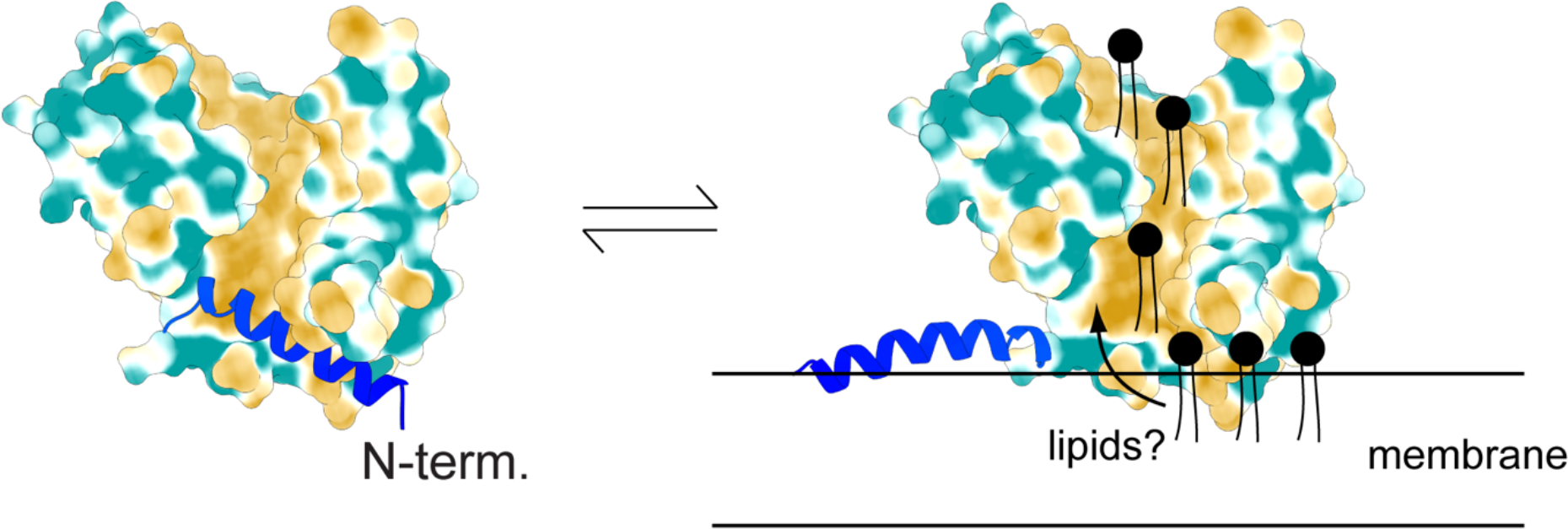
A provisional model of lipid loading and unloading at the N tip. Surface representations of the ATG2A homology model combined with the cartoon representation of the N-terminal helix (blue). If the N-terminal helix undergoes the proposed conformational change upon membrane binding, the hydrophobic cavity of the protein will be readily accessible to lipids on the bound membrane.

## References

AhYoung, A. P., Jiang, J., Zhang, J., Khoi Dang, X., Loo, J. A., Zhou, Z. H., & Egea, P. F. (2015). Conserved SMP domains of the ERMES complex bind phospholipids and mediate tether assembly. Proc Natl Acad Sci U S A, 112(25), E3179–3188. doi:10.1073/pnas.1422363112

Arstila, A. U., & Trump, B. F. (1968). Studies on cellular autophagocytosis. The formation of autophagic vacuoles in the liver after glucagon administration. Am J Pathol, 53(5), 687–733.

Axe, E. L., Walker, S. A., Manifava, M., Chandra, P., Roderick, H. L., Habermann, A., … Ktistakis, N. T. (2008). Autophagosome formation from membrane compartments enriched in phosphatidylinositol 3-phosphate and dynamically connected to the endoplasmic reticulum. J Cell Biol, 182(4), 685–701. doi:10.1083/jcb.200803137

Bakula, D., Muller, A. J., Zuleger, T., Takacs, Z., Franz-Wachtel, M., Thost, A. K., … Proikas-Cezanne, T. (2017). WIPI3 and WIPI4 beta-propellers are scaffolds for LKB1-AMPK-TSC signalling circuits in the control of autophagy. Nat Commun, 8, 15637. doi:10.1038/ncomms15637

Barth, H., Meiling-Wesse, K., Epple, U. D., & Thumm, M. (2001). Autophagy and the cytoplasm to vacuole targeting pathway both require Aut10p. FEBS Lett, 508(1), 23–28.

Baskaran, S., Ragusa, M. J., Boura, E., & Hurley, J. H. (2012). Two-site recognition of phosphatidylinositol 3-phosphate by PROPPINs in autophagy. Mol Cell, 47(3), 339–348. doi:10.1016/j.molcel.2012.05.027

Cheng, J., Fujita, A., Yamamoto, H., Tatematsu, T., Kakuta, S., Obara, K., … Fujimoto, T. (2014). Yeast and mammalian autophagosomes exhibit distinct phosphatidylinositol 3-phosphate asymmetries. Nat Commun, 5, 3207. doi:10.1038/ncomms4207

Choi, A. M., Ryter, S. W., & Levine, B. (2013). Autophagy in human health and disease. N Engl J Med, 368(7), 651–662. doi:10.1056/NEJMra1205406

Chowdhury, S., Otomo, C., Leitner, A., Ohashi, K., Aebersold, R., Lander, G. C., & Otomo, T. (2018). Insights into autophagosome biogenesis from structural and biochemical analyses of the ATG2A-WIPI4 complex. Proc Natl Acad Sci U S A, 115(42), E9792–E9801. doi:10.1073/pnas.1811874115

Dove, S. K., Piper, R. C., McEwen, R. K., Yu, J. W., King, M. C., Hughes, D. C., … Lemmon, M. A. (2004). Svp1p defines a family of phosphatidylinositol 3,5-bisphosphate effectors. EMBO J, 23(9), 1922–1933. doi:10.1038/sj.emboj.7600203

Fujita, N., Hayashi-Nishino, M., Fukumoto, H., Omori, H., Yamamoto, A., Noda, T., & Yoshimori, T. (2008). An Atg4B mutant hampers the lipidation of LC3 paralogues and causes defects in autophagosome closure. Mol Biol Cell, 19(11), 4651–4659. doi:10.1091/mbc.E08-03-0312

Gatta, A. T., & Levine, T. P. (2017). Piecing Together the Patchwork of Contact Sites. Trends Cell Biol, 27(3), 214–229. doi:10.1016/j.tcb.2016.08.010

Geng, J., Baba, M., Nair, U., & Klionsky, D. J. (2008). Quantitative analysis of autophagy-related protein stoichiometry by fluorescence microscopy. J Cell Biol, 182(1), 129–140. doi:10.1083/jcb.200711112

Goddard, T. D., Huang, C. C., Meng, E. C., Pettersen, E. F., Couch, G. S., Morris, J. H., & Ferrin, T. E. (2018). UCSF ChimeraX: Meeting modern challenges in visualization and analysis. Protein Sci, 27(1), 14–25. doi:10.1002/pro.3235

Gomez-Sanchez, R., Rose, J., Guimaraes, R., Mari, M., Papinski, D., Rieter, E., … Reggiori, F. (2018). Atg9 establishes Atg2-dependent contact sites between the endoplasmic reticulum and phagophores. J Cell Biol, 217(8), 2743–2763. doi:10.1083/jcb.201710116

Gopaldass, N., Fauvet, B., Lashuel, H., Roux, A., & Mayer, A. (2017). Membrane scission driven by the PROPPIN Atg18. EMBO J, 36(22), 3274–3291. doi:10.15252/embj.201796859

Graef, M. (2018). Membrane tethering by the autophagy ATG2A-WIPI4 complex. Proc Natl Acad Sci U S A, 115(42), 10540–10541. doi:10.1073/pnas.1814759115

Graef, M., Friedman, J. R., Graham, C., Babu, M., & Nunnari, J. (2013). ER exit sites are physical and functional core autophagosome biogenesis components. Mol Biol Cell, 24(18), 2918–2931. doi:10.1091/mbc.E13-07-0381

Harayama, T., & Riezman, H. (2018). Understanding the diversity of membrane lipid composition. Nat Rev Mol Cell Biol, 19(5), 281–296. doi:10.1038/nrm.2017.138

Hayashi-Nishino, M., Fujita, N., Noda, T., Yamaguchi, A., Yoshimori, T., & Yamamoto, A. (2009). A subdomain of the endoplasmic reticulum forms a cradle for autophagosome formation. Nat Cell Biol, 11(12), 1433–1437. doi:10.1038/ncb1991

Hirata, E., Ohya, Y., & Suzuki, K. (2017). Atg4 plays an important role in efficient expansion of autophagic isolation membranes by cleaving lipidated Atg8 in Saccharomyces cerevisiae. PLoS One, 12(7), e0181047. doi:10.1371/journal.pone.0181047

Itakura, E., & Mizushima, N. (2010). Characterization of autophagosome formation site by a hierarchical analysis of mammalian Atg proteins. Autophagy, 6(6), 764–776. doi:10.4161/auto.6.6.12709

Jeong, H., Park, J., Jun, Y., & Lee, C. (2017). Crystal structures of Mmm1 and Mdm12-Mmm1 reveal mechanistic insight into phospholipid trafficking at ER-mitochondria contact sites. Proc Natl Acad Sci U S A, 114(45), E9502–E9511. doi:10.1073/pnas.1715592114

Kawano, S., Tamura, Y., Kojima, R., Bala, S., Asai, E., Michel, A. H., … Endo, T. (2018). Structure-function insights into direct lipid transfer between membranes by Mmm1-Mdm12 of ERMES. J Cell Biol, 217(3), 959–974. doi:10.1083/jcb.201704119

Kishi-Itakura, C., Koyama-Honda, I., Itakura, E., & Mizushima, N. (2014). Ultrastructural analysis of autophagosome organization using mammalian autophagy-deficient cells. J Cell Sci, 127(Pt 18), 4089–4102. doi:10.1242/jcs.156034

Kotani, T., Kirisako, H., Koizumi, M., Ohsumi, Y., & Nakatogawa, H. (2018). The Atg2-Atg18 complex tethers pre-autophagosomal membranes to the endoplasmic reticulum for autophagosome formation. Proc Natl Acad Sci U S A, 115(41), 10363–10368. doi:10.1073/pnas.1806727115

Kumar, N., Leonzino, M., Hancock-Cerutti, W., Horenkamp, F. A., Li, P., Lees, J. A., … De Camilli, P. (2018). VPS13A and VPS13C are lipid transport proteins differentially localized at ER contact sites. J Cell Biol, 217(10), 3625–3639. doi:10.1083/jcb.201807019

Lamb, C. A., Yoshimori, T., & Tooze, S. A. (2013). The autophagosome: origins unknown, biogenesis complex. Nat Rev Mol Cell Biol, 14(12), 759–774. doi:10.1038/nrm3696

Lang, A. B., John Peter, A. T., Walter, P., & Kornmann, B. (2015). ER-mitochondrial junctions can be bypassed by dominant mutations in the endosomal protein Vps13. J Cell Biol, 210(6), 883–890. doi:10.1083/jcb.201502105

Levine, B., & Kroemer, G. (2019). Biological Functions of Autophagy Genes: A Disease Perspective. Cell, 176(1-2), 11–42. doi:10.1016/j.cell.2018.09.048

Lu, Q., Yang, P., Huang, X., Hu, W., Guo, B., Wu, F., … Zhang, H. (2011). The WD40 repeat PtdIns(3)P-binding protein EPG-6 regulates progression of omegasomes to autophagosomes. Dev Cell, 21(2), 343–357. doi:10.1016/j.devcel.2011.06.024

Meers, P., Ali, S., Erukulla, R., & Janoff, A. S. (2000). Novel inner monolayer fusion assays reveal differential monolayer mixing associated with cation-dependent membrane fusion. Biochim Biophys Acta, 1467(1), 227–243.

Mercer, T. J., Gubas, A., & Tooze, S. A. (2018). A molecular perspective of mammalian autophagosome biogenesis. J Biol Chem, 293(15), 5386–5395. doi:10.1074/jbc.R117.810366

Mizushima, N. (2018). A brief history of autophagy from cell biology to physiology and disease. Nat Cell Biol, 20(5), 521–527. doi:10.1038/s41556-018-0092-5

Mizushima, N., & Komatsu, M. (2011). Autophagy: renovation of cells and tissues. Cell, 147(4), 728–741. doi:10.1016/j.cell.2011.10.026

Mizushima, N., Yamamoto, A., Hatano, M., Kobayashi, Y., Kabeya, Y., Suzuki, K., … Yoshimori, T. (2001). Dissection of autophagosome formation using Apg5-deficient mouse embryonic stem cells. J Cell Biol, 152, 657–668.

Mizushima, N., Yoshimori, T., & Ohsumi, Y. (2011). The role of Atg proteins in autophagosome formation. Annu Rev Cell Dev Biol, 27, 107–132. doi:10.1146/annurev-cellbio-092910-154005

Murley, A., & Nunnari, J. (2016). The Emerging Network of Mitochondria-Organelle Contacts. Mol Cell, 61(5), 648–653. doi:10.1016/j.molcel.2016.01.031

Nguyen, N., Shteyn, V., & Melia, T. J. (2017). Sensing Membrane Curvature in Macroautophagy. J Mol Biol, 429(4), 457–472. doi:10.1016/j.jmb.2017.01.006

Obara, K., Noda, T., Niimi, K., & Ohsumi, Y. (2008). Transport of phosphatidylinositol 3-phosphate into the vacuole via autophagic membranes in Saccharomyces cerevisiae. Genes Cells, 13(6), 537–547. doi:10.1111/j.1365-2443.2008.01188.x

Obara, K., Sekito, T., Niimi, K., & Ohsumi, Y. (2008). The Atg18-Atg2 complex is recruited to autophagic membranes via phosphatidylinositol 3-phosphate and exerts an essential function. Journal of Biological Chemistry.

Otomo, T., Chowdhury, S., & Lander, G. C. (2018). The Rod-Shaped ATG2A-WIPI4 Complex Tethers Membranes In Vitro. Contact, 1, 2515256418819936. doi:10.1177/2515256418819936

Park, J. S., Thorsness, M. K., Policastro, R., McGoldrick, L. L., Hollingsworth, N. M., Thorsness, P. E., & Neiman, A. M. (2016). Yeast Vps13 promotes mitochondrial function and is localized at membrane contact sites. Mol Biol Cell, 27(15), 2435–2449. doi:10.1091/mbc.E16-02-0112

Pfisterer, S. G., Bakula, D., Frickey, T., Cezanne, A., Brigger, D., Tschan, M. P., … Proikas-Cezanne, T. (2014). Lipid droplet and early autophagosomal membrane targeting of Atg2A and Atg14L in human tumor cells. J Lipid Res, 55(7), 1267–1278. doi:10.1194/jlr.M046359

Polson, H. E., de Lartigue, J., Rigden, D. J., Reedijk, M., Urbe, S., Clague, M. J., & Tooze, S. A. (2010). Mammalian Atg18 (WIPI2) localizes to omegasome-anchored phagophores and positively regulates LC3 lipidation. Autophagy, 6(4), 506–522. doi:10.4161/auto.6.4.11863

Polson, H. E. J., Girardin, S. E., Wilson, M. I., & Tooze, S. A. (2014). WIPI2 links LC3 conjugation with PI3P, autophagosome formation, and pathogen clearance by recruiting Atg12–5-16L1. Molecular cell.

Proikas-Cezanne, T., Takacs, Z., Donnes, P., & Kohlbacher, O. (2015). WIPI proteins: essential PtdIns3P effectors at the nascent autophagosome. J Cell Sci, 128(2), 207–217. doi:10.1242/jcs.146258

Proikas-Cezanne, T., Waddell, S., Gaugel, A., Frickey, T., Lupas, A., & Nordheim, A. (2004). WIPI-1alpha (WIPI49), a member of the novel 7-bladed WIPI protein family, is aberrantly expressed in human cancer and is linked to starvation-induced autophagy. Oncogene, 23(58), 9314–9325. doi:10.1038/sj.onc.1208331

Reggiori, F., & Klionsky, D. J. (2013). Autophagic processes in yeast: mechanism, machinery and regulation. Genetics, 194(2), 341–361. doi:10.1534/genetics.112.149013

Roy, A., Kucukural, A., & Zhang, Y. (2010). I-TASSER: a unified platform for automated protein structure and function prediction. Nat Protoc, 5(4), 725–738. doi:10.1038/nprot.2010.5

Scacioc, A., Schmidt, C., Hofmann, T., Urlaub, H., Kuhnel, K., & Perez-Lara, A. (2017). Structure based biophysical characterization of the PROPPIN Atg18 shows Atg18 oligomerization upon membrane binding. Sci Rep, 7(1), 14008. doi:10.1038/s41598-017-14337-5

Schauder, C. M., Wu, X., Saheki, Y., Narayanaswamy, P., Torta, F., Wenk, M. R., … Reinisch, K. M. (2014). Structure of a lipid-bound extended synaptotagmin indicates a role in lipid transfer. Nature, 510(7506), 552–555. doi:10.1038/nature13269

Shintani, T., Suzuki, K., Kamada, Y., Noda, T., & Ohsumi, Y. (2001). Apg2p functions in autophagosome formation on the perivacuolar structure. J Biol Chem, 276(32), 30452–30460. doi:10.1074/jbc.M102346200

Struck, D. K., Hoekstra, D., & Pagano, R. E. (1981). Use of resonance energy transfer to monitor membrane fusion. Biochemistry, 20(14), 4093–4099.

Suzuki, K., Akioka, M., Kondo-Kakuta, C., Yamamoto, H., & Ohsumi, Y. (2013). Fine mapping of autophagy-related proteins during autophagosome formation in Saccharomyces cerevisiae. J Cell Sci, 126(Pt 11), 2534–2544. doi:10.1242/jcs.122960

Taguchi-Atarashi, N., Hamasaki, M., Matsunaga, K., Omori, H., Ktistakis, N. T., Yoshimori, T., & Noda, T. (2010). Modulation of local PtdIns3P levels by the PI phosphatase MTMR3 regulates constitutive autophagy. Traffic, 11(4), 468–478. doi:10.1111/j.1600-0854.2010.01034.x

Tamura, N., Nishimura, T., Sakamaki, Y., Koyama-Honda, I., Yamamoto, H., & Mizushima, N. (2017). Differential requirement for ATG2A domains for localization to autophagic membranes and lipid droplets. FEBS Lett, 591(23), 3819–3830. doi:10.1002/1873-3468.12901

Tooze, S. A., & Yoshimori, T. (2010). The origin of the autophagosomal membrane. Nat Cell Biol, 12(9), 831–835. doi:10.1038/ncb0910-831

Uemura, T., Yamamoto, M., Kametaka, A., Sou, Y. S., Yabashi, A., Yamada, A., … Waguri, S. (2014). A cluster of thin tubular structures mediates transformation of the endoplasmic reticulum to autophagic isolation membrane. Mol Cell Biol, 34(9), 1695–1706. doi:10.1128/MCB.01327-13

Velayos-Baeza, A., Vettori, A., Copley, R. R., Dobson-Stone, C., & Monaco, A. P. (2004). Analysis of the human VPS13 gene family. Genomics, 84(3), 536–549. doi:10.1016/j.ygeno.2004.04.012

Velikkakath, A. K., Nishimura, T., Oita, E., Ishihara, N., & Mizushima, N. (2012). Mammalian Atg2 proteins are essential for autophagosome formation and important for regulation of size and distribution of lipid droplets. Mol Biol Cell, 23(5), 896–909. doi:10.1091/mbc.E11-09-0785

Wang, C. W., Kim, J., Huang, W. P., Abeliovich, H., Stromhaug, P. E., Dunn, W. A., Jr., & Klionsky, D. J. (2001). Apg2 is a novel protein required for the cytoplasm to vacuole targeting, autophagy, and pexophagy pathways. J Biol Chem, 276(32), 30442–30451. doi:10.1074/jbc.M102342200

Wong, L. H., Gatta, A. T., & Levine, T. P. (2018). Lipid transfer proteins: the lipid commute via shuttles, bridges and tubes. Nat Rev Mol Cell Biol. doi:10.1038/s41580-018-0071-5

Yamamoto, A., Masaki, R., Fukui, Y., & Tashiro, Y. (1990). Absence of cytochrome P-450 and presence of autolysosomal membrane antigens on the isolation membranes and autophagosomal membranes in rat hepatocytes. J Histochem Cytochem, 38(11), 1571–1581. doi:10.1177/38.11.2212617

Yla-Anttila, P., Vihinen, H., Jokitalo, E., & Eskelinen, E. L. (2009). 3D tomography reveals connections between the phagophore and endoplasmic reticulum. Autophagy, 5(8), 1180–1185. doi:10274 [pii]

Zheng, J. X., Li, Y., Ding, Y. H., Liu, J. J., Zhang, M. J., Dong, M. Q., … Yu, L. (2017). Architecture of the ATG2B-WDR45 complex and an aromatic Y/HF motif crucial for complex formation. Autophagy, 13(11), 1870–1883. doi:10.1080/15548627.2017.1359381

